# Remodeling of Chemotaxis is a Cornerstone of Bacterial Swarming

**DOI:** 10.1101/2020.04.27.064519

**Authors:** Jonathan D. Partridge, Nhu Q. Nguyen, Yann S. Dufour, Rasika M. Harshey

## Abstract

Many bacteria use flagella-driven motility to swarm or move collectively over a surface terrain. Bacterial adaptations for swarming can include cell elongation, hyper-flagellation, recruitment of special stator proteins and surfactant secretion, among others. We recently demonstrated another swarming adaptation in *Escherichia coli*, wherein the chemotaxis pathway is remodeled to increase run durations (decrease tumble bias), with running speeds increased as well. We show here that the modification of motility parameters during swarming is not unique to *E. coli*, but shared by a diverse group of bacteria we examined – *Proteus mirabilis, Serratia marcescens, Salmonella enterica, Bacillus subtilis*, and *Pseudomonas aeruginosa* – suggesting that altering the chemosensory physiology is a cornerstone of swarming.

**Importance:** Bacteria within a swarm move characteristically in packs, displaying an intricate swirling motion where hundreds of dynamic packs continuously form and dissociate as the swarm colonizes increasing expanse of territory. The demonstrated property of *E. coli* to reduce its tumble bias and hence increase its run duration during swarming is expected to maintain/promote side-by-side alignment and cohesion within the bacterial packs. Here we observe a similar low tumble bias in five different bacterial species, both Gram positive and Gram negative, each inhabiting a unique habitat and posing unique problems to our health. The unanimous display of an altered run-tumble bias in swarms of all species examined here suggests that this behavioral adaptation is crucial for swarming.

## Introduction

Swarming is defined as a rapid collective migration of bacteria across a surface, powered by flagella (1-3). A wide array of phenotypic adaptations are associated with swarming. A common attribute of all swarms is a pattern of ceaseless circling motion, in which packs of cells all traveling in the same directions split and merge, with continuous exchange of bacteria between the packs (3-5). This behavior differs from movement of the bacteria in bulk liquid, where they swim individually (6). In *E. coli*, the mechanics of flagella are similar during both swimming and swarming in that peritrichous flagella driven by bi-directional rotary motors switch between counterclockwise (CCW) and clockwise (CW) directions. However, while CCW rotation promotes formation of a coherent flagellar bundle that propels the cell forward (run) during both swimming and swarming, a transient switch in rotational direction (CW) causes the cell to tumble while swimming, but reverse direction while swarming (7, 8).

The switching frequency of the flagellar motor is controlled by the chemotaxis system, best studied in *E. coli*, where transmembrane receptors detect extracellular signals and transmit them via phosphorelay to the motor, to promote migration to favorable locales during swimming (9). The ability to perform chemotaxis is not essential for swarming, but a basal tumble bias is important (10). We recently reported that compared to planktonic cells, *E. coli* taken from a swarm exhibit more highly extended runs and higher speeds, and that this low tumble bias displayed by swarmers is the optimal bias for maximizing swarm expansion (11). Post-transcriptional changes that alter the levels of a key signaling protein suggested that the chemotaxis signaling pathway is reprogrammed for swarming. A low tumble bias (TB) is consistent with the superdiffusive Lévy walk run trajectories observed in swarms of *S. marcescens* and *B. subtilis* (12), and could improve swarming performance at the minimum by favoring the alignment of cells all travelling in the same direction in a pack. Whether a low TB facilitates expansion of the swarm by improving chemotactic performance is not known, but a functional chemotaxis system is apparently necessary for swarmers to avoid antibiotics (13). Swarming allows bacteria opportunities for dispersal in ecological niches and contributes to pathogenicity in many species (14), notably in conferring enhanced resistance to antibiotics (13).

Here, we examined TB and speeds during swarming in a selected mix of swarmer species, united only in their macroscopic display of swirling packs. *P. mirabilis* will swarm on hard agar (1.5% agar and above; ‘robust’ conditions), but all other species will only swarm on softer agar (0.5% to 0.8% agar; ‘temperate’ conditions). *P. aeruginosa* has a polar flagellum (15), while the others are all peritrichously flagellated. Except for *S. enterica*, swarming is aided by secretion of surfactants or polysaccharides in the rest. *P. mirabilis* can elongate substantially (10-80 μm) on hard agar (16), while the others do not change morphology dramatically. Despite these varying swarming adaptations, we find that they all share the same low TB and higher run speeds as first reported for *E. coli*, suggesting that this behavior is a universal adaptation for successful migration on a surface.

## Results and Discussion

The methodology and growth conditions used to monitor TB and speed in this study were similar to those used for *E. coli* (11), and were consistently applied across all swarming species (see SI). Swarming was first described in *Proteus* species in 1885 (17). Temperate swarming conditions were first identified in *S. marcescens* (18), followed by in *E. coli* and *S. enterica* (19), as well as in a large number or other species (see Fig. 1 in (20)), including in *B. subtilis* (21), and *P. aeruginosa* (22). To maintain uniformity in tumble behavior, we bypassed some swarm-related phenotypes of individual species. For example, *P. mirabilis* gets extremely long on hard agar, and long cells will not tumble. Under temperate conditions used here, their cell length (2.5 ± 0.7 μm, *n* = 50) was unchanged from those cultivated in liquid (2.1 ± 0.5 μm, *n* = 50). *S. marcescens* secretes serrawettin, a cyclic lipopeptide surfactant (3). Preliminary tracking experiments with *S. marcescens* cells taken from liquid showed large circular trajectories (Fig. S1, left). Such trajectories have been observed with *E. coli* and *Caulobacter crescentus* when swimming close to a glass surface (23). We therefore used an *S. marcescens* mutant deficient in serrawettin production (Fig. S1, right), which abolished the circular motion. *B. subtilis* makes a similar surfactant (2), so we used a *srfA* mutant deficient in surfactin synthesis. *P. aeruginosa* motility in liquid differs from the run-tumble pattern, and is instead characterized as a run-reverse-turn pattern, where prolonged runs are interrupted by a reversal and ‘flick’ to cause a change in direction (24). The tumble angle distribution plots we observed were consistent with run-reverse-flick. While technically *P. aeruginosa* does not tumble, in our analysis, the run-reverse and reverse-flick are both identified as tumbles. We will discuss our findings in the order of discovery of swarming in the bacterial species studied here.

**Figure 1.**
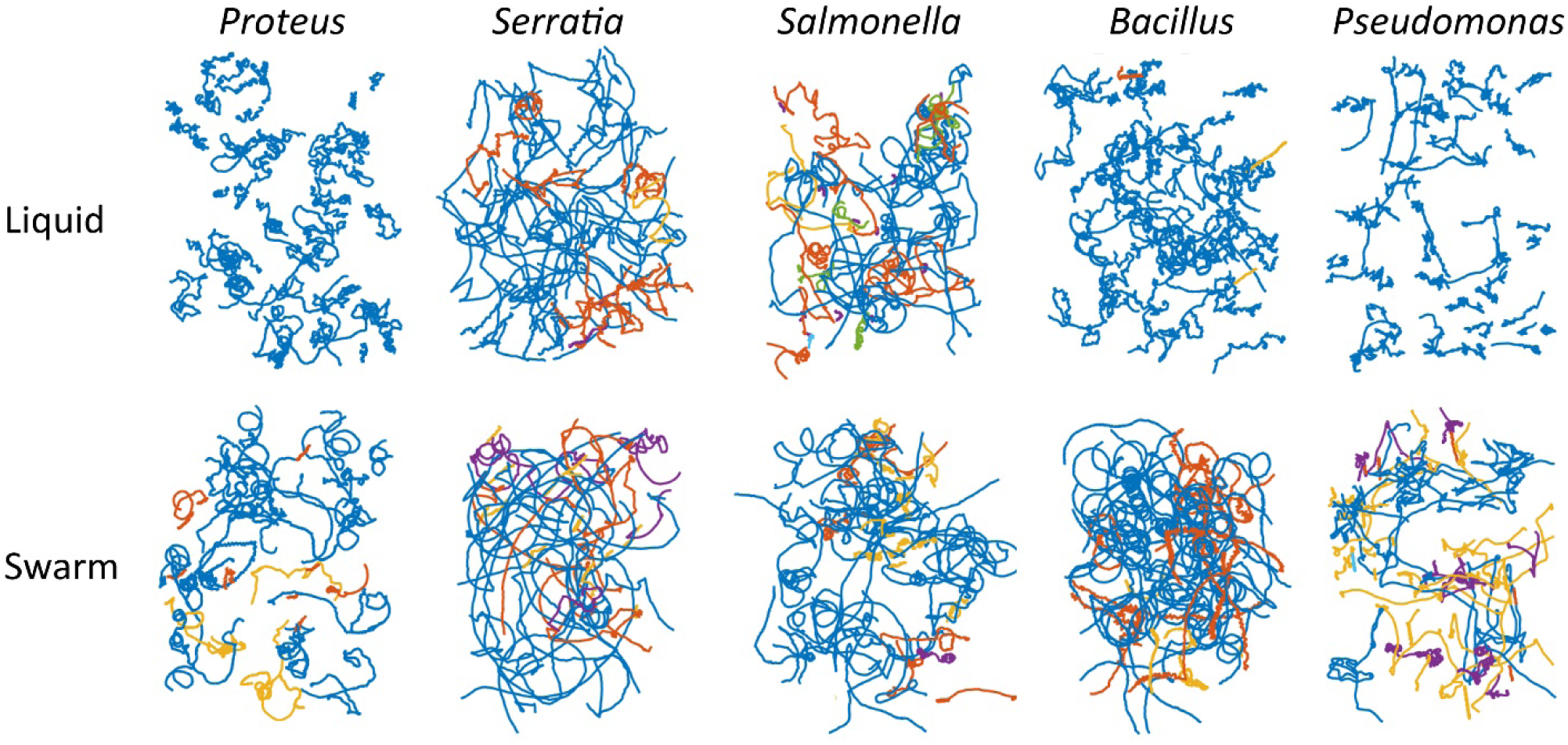
Trajectories of *Proteus, Serratia, Salmonella, Bacillus*, and *Proteus* cells cultivated in liquid or swarm conditions. Cells were grown in LB (liquid) or LB swarm agar, each supplemented with glucose (0.5 % w/v), before transfer to LB liquid for observation in a pseudo-2D environment. Cell movement was recorded for 100 s using phase-contrast microscopy at 10X magnification. Trajectories of single representative experiments shown. Different colors correspond to individual tracks.

Representative cell trajectories in liquid or swarm media for all bacterial species tested are shown in Figure 1. All show a distinct shift in motion paths under the two conditions, becoming smoother (long run trajectories) during swarming. Quantitative analyses of these trajectories are shown in Figure 2. The changes in median TB values from liquid to swarm are as follows. *P. mirabilis*: 0.27 to 0.14, *S. marcescens*: 0.23 to 0.037, *S. enterica*: 0.07 to 0.05, *B. subtilis*: 0.24 to 0.048. *P. aeruginosa*: 0.53 to 0.31 (stats. detailed in Table S1). While the overall pattern was that TBs shifted to lower values during swarming, we note that TB values for *S. enterica* are lower than *E. coli* in liquid to begin with, as reported in single motor assays (25). For comparison, TB values for *E. coli* decreased from a median of 0.12 in liquid, to 0.04 in swarmers (11).

**Figure 2.**
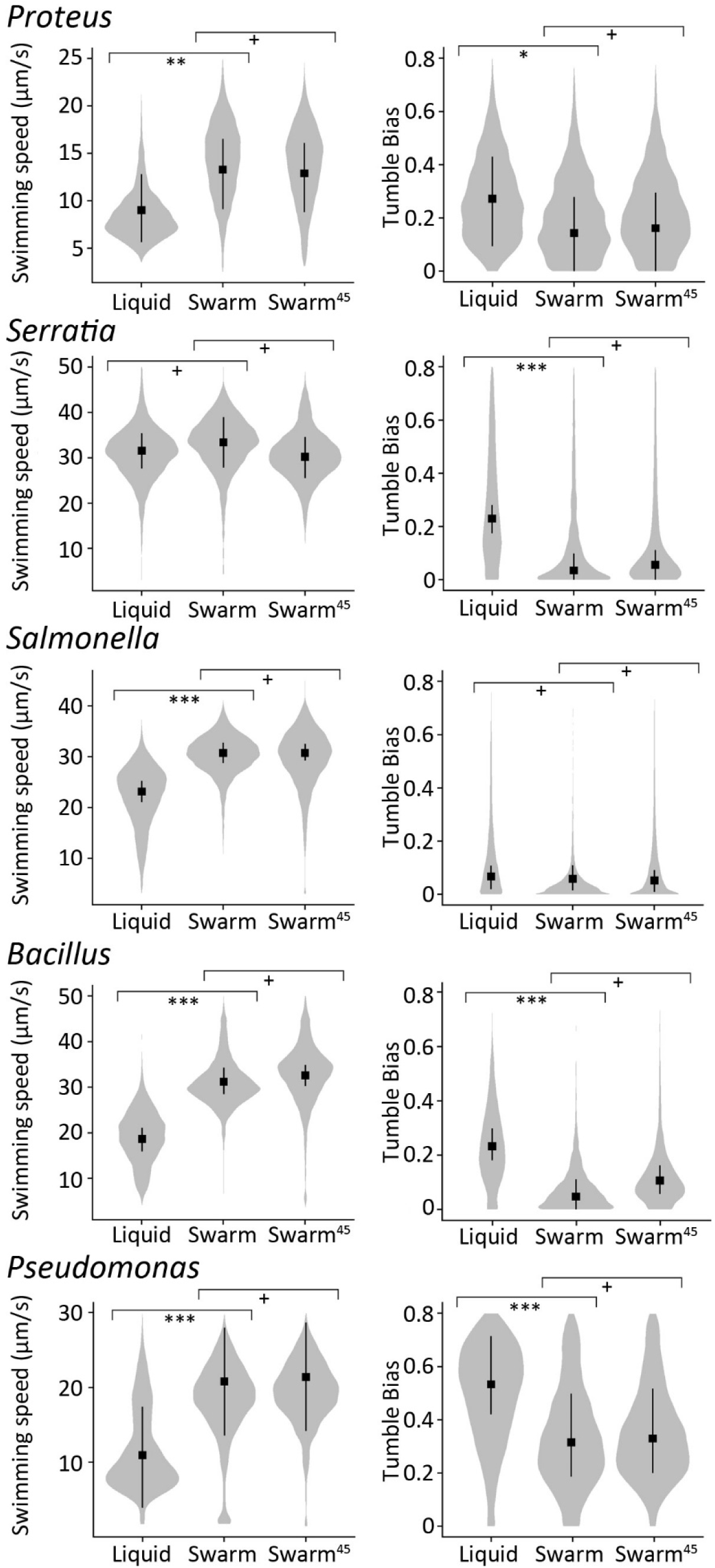
Swimming speed and tumble bias of *Proteus, Serratia, Salmonella, Bacillus*, and *Pseudomonas* cells cultivated in liquid, swarm, or swarm^45^ conditions. Cells were grown in LB (liquid) or LB swarm agar, each supplemented with glucose (0.5 % w/v), before transfer to LB liquid for observation in a pseudo-2D environment. Swarm^45^ denotes isolated ‘swarm’ samples monitored again after 45 min had elapsed. Cell movement was recorded for 100 s using phase-contrast microscopy at 10X magnification. Probability distribution of swimming speeds (micrometers per second) (left) and cell tumble biases (right) shown. Distribution of each parameter was calculated from more than 4600 individual trajectories (> 1000 min of cumulative time) for each condition, from at least three independent experiments. The square and bars indicates the mean and 95% credible intervals of the posterior probabilities of the medians for each treatment. Calculated *P* values are indicated: *, <0.05, **, <0.01, or ***, <0.0001. +, *P* value >0.05.

The low TB displayed by *E. coli* swarmers was observed to be stable up to 45 minutes, and persisted through one cell division at room temperature (∼120 minutes) (11). We therefore also included a 45-minute time point (after lifting cells from the swarm) for tracking all five swarmers. At 45 minutes post-removal from the swarm, most bacteria maintained their low TB values (stats. found in Table S1).

As observed for *E. coli*, running speeds (μm/sec) for a majority of the bacterial species increased significantly between liquid and swarm as follows. *P. mirabilis*: 9.01 to 13.3, *S. enterica*: 23.1 to 30.7, *B. subtilis*: 18.6 to 31, *P. aeruginosa*: 21.9 to 41.6 (stats. in Table S1). These values for *E. coli* were 21 µm/sec in liquid, and 25 µm/s in swarmers (11).

## Summary

Keeping swarming conditions the same, we demonstrate here that despite different natural habitats and widely different swarming adaptations discovered in the laboratory, the swarmers studied here all modify their TB, and a majority modify run speeds during swarming, similar to that reported for *E. coli* (11). This apparently common behavior suggests that it represents a successful strategy for collective migration across a surface. In *E. coli*, elevation or stabilization of the chemotaxis component CheZ was shown to be responsible for the low TB (11). The higher motor torque and speed recorded for single motors of swarmers likely represent increased proton-motive force resulting from the altered swarmer physiology. For example, *S. enterica* swarmers are reported to upregulate tricarboxylic acid (TCA) cycle enzymes (26) and swarming patterns in *Proteus* are contingent on a complete TCA cycle (27). Similar changes in metabolism may fuel the increased speeds in the other bacteria. Future work will reveal the mechanisms used by each of these bacteria to arrive at what is apparently a common solution for maximizing collective motion.

## Materials and methods

Strains used in this study are described in Table S1. Cell culture and swarm setup are described in supplementary materials. Tracking experiments and analysis were largely carried out as described previously (11). For details and changes, see supplementary materials

## Acknowledgments

J.D.P. and R.M.H. conceptualized the study. J.D.P. and N.Q.N. performed the experiments. J.D.P., Y.S.D. and R.M.H. analyzed the data and wrote the manuscript.

## Supplementary Figure Legends

**Figure S1.**
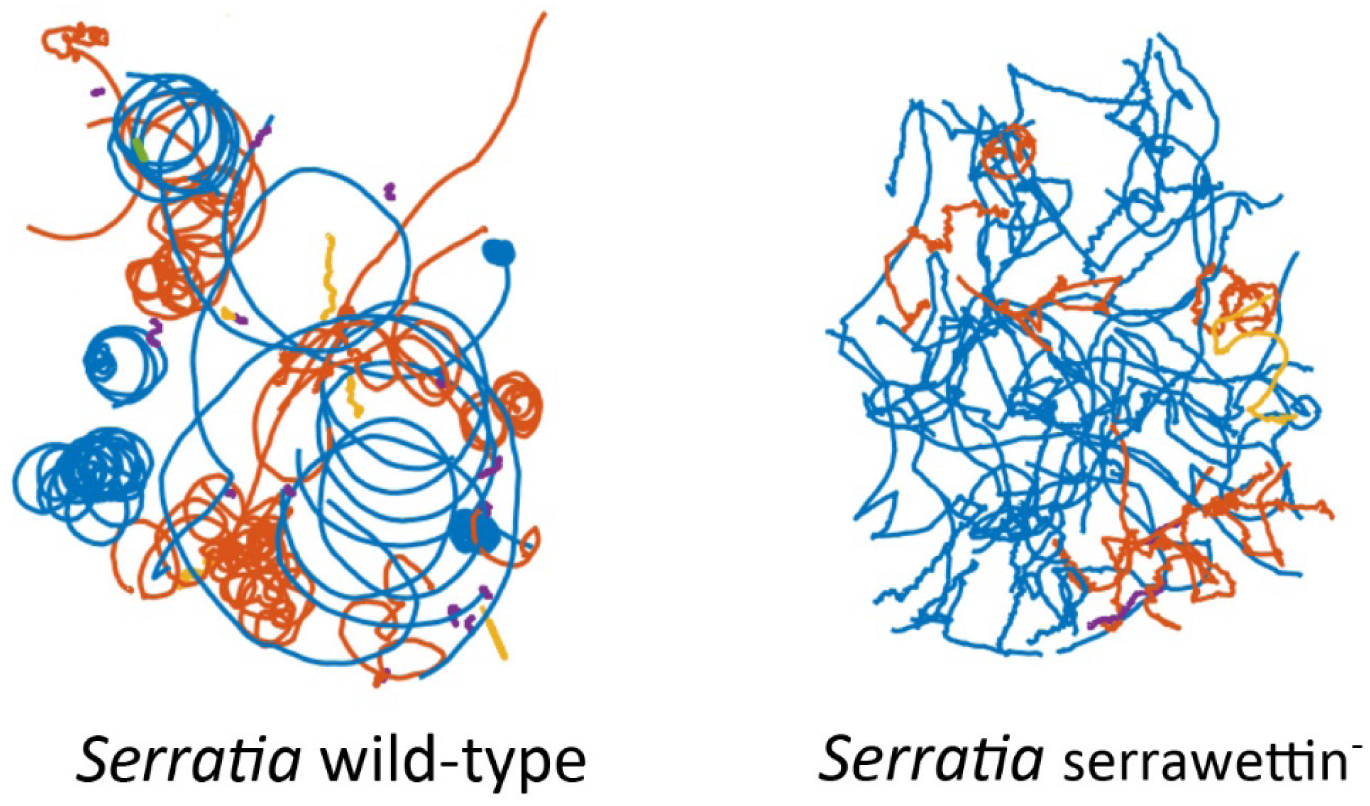
Representative trajectories of wild-type *Serratia marcescens* and RH1041, a serrawettin-mutant. Cells were grown in LB (liquid) plus glucose (0.5% w/v), before transfer to LB liquid for observation. Cell movement was recorded for 100 s using phase-contrast microscopy at 10X magnification. Trajectories of single representative experiments shown. Different colors correspond to individual tracks.

## Supplementary Material

### Strains and growth conditions

Strains used in this study were: *Salmonella enterica* 14028 and *Serratia marcescens* 274 were sourced from the American Type Culture Collection, *S. marcescens* serrawettin^-^ (RH1041, SMu4e in (1)), *Bacillus subtilis srfA* (DS191; gift from Daniel Kearns), *Pseudomonas aeruginosa* (PA01; gift from Verinita Gordon), and *Proteus mirabilis* (lab collection). Cells were cultured in Lennox Broth (LB, 10 g/L tryptone, 5g/L yeast extract, 5g/L NaCl). Starting from single colonies isolated on agar plates, cells were grown overnight in broth cultures and sub-cultured using 1:100 dilution ratio in fresh medium and grown for around 4 h to an optical density at 600 nm (OD_600_) of 0.4. Liquid cultures were grown at 30°C in an Erlenmeyer flask on an orbital shaker at 200 r.p.m. for aeration. Swarm plates (LB solidified with 0.5 % Eiken agar [Eiken Chemical Co., Japan], respectively) were poured and held at room temperature for 16 hours prior to inoculation with 6 µl of an overnight culture in the center and incubated at 30°C. All media was supplemented with 0.5% glucose. For experiments with edge cells in a swarm, cells were collected after 4 hours by gently washing the cells from the edge and resuspended in LB glucose for tracking assays (see below).

### Time-lapse microscopy, cell tracking, and trajectory analysis

Cells were harvested (2,000 g, 5 minutes) and washed twice in fresh media. They were tracked at room temperature in LB supplemented with 0.5% glucose (w/v). Resuspended cells were diluted to an OD_600_ of ∼ 0.01-0.05, and 5 μl were introduced between a glass microscope slide and 22 mm^2^ #1.5 coverslip, sealed using nail varnish. This created a channel ∼10 μm deep. Swimming cells were recorded at 10 frames per second with a Olympus XM10 camera (1,376 x 1,032 pixels, 10 ms exposure) mounted on an inverted microscope (Olympus BX53) with a 10X phase contrast objective (Olympus PLN 10X). The field of view was ∼0.9 mm square containing on average 200 to 600 cells. Cell trajectories were reconstructed using a custom MATLAB (Mathworks) code (github.com/dufourya/SwimTracker) (2, 3). Behavioral parameters such as speed and tumble bias, were extracted from single-cell trajectories as previously described (2). The swimming speed was calculated by taking the average velocity of individual cells over their respective trajectories excluding the frames where cells are predicted to be tumbling. Trajectories shorter than 5 seconds were discarded. Cells with a diffusion coefficient of less than 10 m^2^/s are driven only by Brownian motion and were classified as non-motile and not included in the analyses of swimming speed and tumble bias.

Bayesian sampling was used to determine if the medians of the swimming speed and tumble bias are significantly different between liquid, swarm, and swarm^45^ preparations. The posterior probability distributions of the medians for each strain and each treatment were calculated using a linear mixed-effect model ((Swimming_speed, Tumble_bias) ∼ Treatment + 1|Replicate) with a Gaussian distribution link function. Each cell trajectory was weighted by its length to obtain an accurate quantification of cell-to-cell variability in the population. All the statistical analyses were done by sampling of the respective mixed-effect generalized linear models using the RSTAN (5) and BRMS packages (6) in R (7) with 4 chains, each with 1,000 warmup iterations and at least 5,000 sampling iterations. P values and credible intervals were calculated by sampling the posterior probability distributions. Uninformative priors were set to the defaults generated by BRMS. The plots were generated using the ggplot2 (8) and tidybayes (9) packages.

---

**Table S1.** Mean posterior probabilities for the median tumble biases and swimming speeds and their comparisons of *Proteus, Serratia, Salmonella, Bacillus*, and *Pseudomonas* cells cultivated in liquid, swarm, or swarm^45^ conditions. Bayesian sampling was used to determine if the medians of the swimming speed and tumble bias are significantly different between liquid, swarm, and swarm^45^ preparations. Swarm^45^ denotes isolated ‘swarm’ samples monitored again after 45 min had elapsed. The posterior probability distributions of the medians for each strain and each treatment were calculated using a linear mixed-effect model with a Gaussian distribution link function. The mean and 95% credible intervals^a^ of the posteriors of the medians for each distribution is also reported. See supplementary information for more details. The means and 95% credible intervals^b^ of the differences of the medians between conditions is reported. *P* values^c^ (for difference in the medians >0 or <0) were calculated by sampling the posterior probability distributions.

|  | Median posteriors<sup>a</sup> [95% CI] |  | Conditions contrast<sup>b</sup> [95% CI] | P value<sup>c</sup> |
| --- | --- | --- | --- | --- |
| ***Proteus* Speeds** |  |  |  |  |
| Liquid | 9.01 [5.63, 12.81] | Liquid vs. Swarm | -4.28 [-7.55, -1.59] | 2.4E-3 |
| Swarm | 13.29 [9.10, 16.52] | Swarm vs. Swarm<sup>45</sup> | -0.39 [-1.88, 1.11] | 2.2E-1 |
| Swarm<sup>45</sup> | 12.90 [8.78, 16.10] | Liquid vs. Swarm<sup>45</sup> | -3.89 [-7.13, -1.19] | 4.1E-3 |
| ***Proteus* TB** |  |  |  |  |
| Liquid | 0.27 [0.09, 0.42] | Liquid vs. Swarm | 0.13 [-0.01, 0.27] | 3.1E-2 |
| Swarm | 0.14 [0, 0.28] | Swarm vs. Swarm<sup>45</sup> | 0.02 [-0.05, 0.09] | 2.4E-1 |
| Swarm<sup>45</sup> | 0.16 [0, 0.29] | Liquid vs. Swarm<sup>45</sup> | 0.11 [-0.02, 0.25] | 4.6E-2 |
| ***Serratia* Speeds** |  |  |  |  |
| Liquid | 31.56 [27.59, 35.37] | Liquid vs. Swarm | -1.85 [-7.16, 3.46] | 2.3E-1 |
| Swarm | 33.41 [27.86, 38.98] | Swarm vs. Swarm<sup>45</sup> | -3.22 [-8.75, 2.51] | 1.2E-1 |
| Swarm<sup>45</sup> | 30.18 [25.50, 34.60] | Liquid vs. Swarm<sup>45</sup> | 1.37 [-2.89, 5.61] | 2.4E-1 |
| ***Serratia* TB** |  |  |  |  |
| Liquid | 0.23 [0.17, 0.28] | Liquid vs. Swarm | 0.19 [0.14, 0.25] | <1E-05 |
| Swarm | 0.04 [0, 0.10] | Swarm vs. Swarm<sup>45</sup> | 0.02 [-0.03, 0.08] | 2.0E-1 |
| Swarm<sup>45</sup> | 0.06 [0, 0.11] | Liquid vs. Swarm<sup>45</sup> | 0.19 [0.14, 0.25] | <1E-05 |
| ***Salmonella* Speeds** |  |  |  |  |
| Liquid | 23.12 [21.02, 25.19] | Liquid vs. Swarm | -7.59 [-9.84, -5.36] | <1E-05 |
| Swarm | 30.71 [29.24, 32.47] | Swarm vs. Swarm<sup>45</sup> | -0.01 [-2.41, 2.26] | 5.0E-1 |
| Swarm<sup>45</sup> | 30.70 [28.75, 32.72] | Liquid vs. Swarm<sup>45</sup> | -7.58 [-10.39, -4.84] | 1.4E-04 |
| ***Salmonella* TB** |  |  |  |  |
| Liquid | 0.07 [0.02, 0.11] | Liquid vs. Swarm | 0.02 [-0.00, 0.04] | 6.7E-2 |
| Swarm | 0.05 [0.01, 0.09] | Swarm vs. Swarm<sup>45</sup> | 0.01 [-0.04, 0.037] | 3.1E-1 |
| Swarm<sup>45</sup> | 0.06 [0.02, 0.11] | Liquid vs. Swarm<sup>45</sup> | 0.01 [-0.03, 0.07] | 4.1E-1 |
| ***Bacillus* Speeds** |  |  |  |  |
| Liquid | 18.64 [15.87, 21.10] | Liquid vs. Swarm | -12.57 [-15.87, -8.83] | <1E-05 |
| Swarm | 31.20 [28.43, 34.31] | Swarm vs. Swarm<sup>45</sup> | 1.37 [-1.37, 4.78] | 2.1E-1 |
| Swarm<sup>45</sup> | 32.58 [30.26, 34.86] | Liquid vs. Swarm<sup>45</sup> | -13.94 [-15.74, -11.69] | <1E-05 |
| ***Bacillus* TB** |  |  |  |  |
| Liquid | 0.23 [0.18, 0.30] | Liquid vs. Swarm | 0.18 [0.11, 0.27] | 2.5E-4 |
| Swarm | 0.048 [0, 0.11] | Swarm vs. Swarm<sup>45</sup> | 0.06 [-0.02, 0.13] | 5.0E-2 |
| Swarm<sup>45</sup> | 0.11 [0.06, 0.16] | Liquid vs. Swarm<sup>45</sup> | 0.13 [0.07, 0.18] | 6.5E-4 |
| ***Pseudomonas* Speeds** |  |  |  |  |
| Liquid | 21.91 [7.92, 34.90] | Liquid vs. Swarm | -19.70 [-28.33, -9.76] | 9.0E-04 |
| Swarm | 41.60 [27.10, 55.96] | Swarm vs. Swarm<sup>45</sup> | 1.14 [-8.66, 11.02] | 4.0E-1 |
| Swarm<sup>45</sup> | 42.75 [28.41, 57.34] | Liquid vs. Swarm<sup>45</sup> | -20.83 [-29.60, -10.58] | 7.0E-4 |
| ***Pseudomonas* TB** |  |  |  |  |
| Liquid | 0.53 [0.42, 0.71] | Liquid vs. Swarm | 0.22 [0.13, 0.30] | 3.0E-4 |
| Swarm | 0.31 [0.19, 0.50] | Swarm vs. Swarm<sup>45</sup> | 0.015 [-0.08, 0.11] | 3.7E-1 |
| Swarm<sup>45</sup> | 0.33 [0.20, 0.52] | Liquid vs. Swarm<sup>45</sup> | 0.20 [0.11, 0.29] | 1.0E-3 |

## References

1. Harshey RM. 2003. Bacterial motility on a surface: many ways to a common goal. Annu Rev Microbiol 57:249–73.

2. Kearns DB. 2010. A field guide to bacterial swarming motility. Nat Rev Microbiol 8:634–44.

3. Partridge JD, Harshey RM. 2013. Swarming: flexible roaming plans. J Bacteriol 195:909–18.

4. Be’er A, Ariel G. 2019. A statistical physics view of swarming bacteria. Mov Ecol 7:9.

5. Darnton NC, Turner L, Rojevsky S, Berg HC. 2010. Dynamics of bacterial swarming. Biophys J 98:2082–90.

6. Berg HC. 2003. The rotary motor of bacterial flagella. Annu Rev Biochem 72:19–54.

7. Turner L, Ryu WS, Berg HC. 2000. Real-time imaging of fluorescent flagellar filaments. J Bacteriol 182:2793–801.

8. Turner L, Zhang R, Darnton NC, Berg HC. 2010. Visualization of Flagella during bacterial Swarming. J Bacteriol 192:3259–67.

9. Parkinson JS, Hazelbauer GL, Falke JJ. 2015. Signaling and sensory adaptation in Escherichia coli chemoreceptors: 2015 update. Trends Microbiol 23:257–66.

10. Mariconda S, Wang Q, Harshey RM. 2006. A mechanical role for the chemotaxis system in swarming motility. Mol Microbiol 60:1590–602.

11. Partridge JD, Nhu NTQ, Dufour YS, Harshey RM. 2019. Escherichia coli Remodels the Chemotaxis Pathway for Swarming. mBio 10.

12. Ariel G, Rabani A, Benisty S, Partridge JD, Harshey RM, Be’er A. 2015. Swarming bacteria migrate by Levy Walk. Nat Commun 6:8396.

13. Partridge JD, Ariel G, Schvartz O, Harshey RM, Be’er A. 2018. The 3D architecture of a bacterial swarm has implications for antibiotic tolerance. Sci Rep 8:15823.

14. Duan Q, Zhou M, Zhu L, Zhu G. 2013. Flagella and bacterial pathogenicity. J Basic Microbiol 53:1–8.

15. Kohler T, Curty LK, Barja F, van Delden C, Pechere JC. 2000. Swarming of Pseudomonas aeruginosa is dependent on cell-to-cell signaling and requires flagella and pili. J Bacteriol 182:5990–6.

16. Belas R, Erskine D, Flaherty D. 1991. Proteus mirabilis mutants defective in swarmer cell differentiation and multicellular behavior. J Bacteriol 173:6279–88.

17. Henrichsen J. 1972. Bacterial surface translocation: a survey and a classification. Bacteriol Rev 36:478–503.

18. Alberti L, Harshey RM. 1990. Differentiation of Serratia marcescens 274 into swimmer and swarmer cells. J Bacteriol 172:4322–8.

19. Harshey RM, Matsuyama T. 1994. Dimorphic transition in Escherichia coli and Salmonella typhimurium: surface-induced differentiation into hyperflagellate swarmer cells. Proc Natl Acad Sci U S A 91:8631–5.

20. Harshey RM. 1994. Bees aren’t the only ones: swarming in gram-negative bacteria. Mol Microbiol 13:389–94.

21. Kearns DB, Losick R. 2003. Swarming motility in undomesticated Bacillus subtilis. Mol Microbiol 49:581–90.

22. Rashid MH, Kornberg A. 2000. Inorganic polyphosphate is needed for swimming, swarming, and twitching motilities of Pseudomonas aeruginosa. Proc Natl Acad Sci U S A 97:4885–90.

23. Lauga E, DiLuzio WR, Whitesides GM, Stone HA. 2006. Swimming in circles: motion of bacteria near solid boundaries. Biophys J 90:400–12.

24. Qian C, Wong CC, Swarup S, Chiam KH. 2013. Bacterial tethering analysis reveals a “run-reverse-turn” mechanism for Pseudomonas species motility. Appl Environ Microbiol 79:4734–43.

25. Partridge JD, Nieto V, Harshey RM. 2015. A new player at the flagellar motor: FliL controls both motor output and bias. mBio 6:e02367.

26. Kim W, Surette MG. 2004. Metabolic differentiation in actively swarming Salmonella. Mol Microbiol 54:702–14.

27. Alteri CJ, Himpsl SD, Engstrom MD, Mobley HL. 2012. Anaerobic respiration using a complete oxidative TCA cycle drives multicellular swarming in Proteus mirabilis. mBio 3.

## References

1. Matsuyama T, Bhasin A, Harshey RM. 1995. Mutational analysis of flagellum-independent surface spreading of Serratia marcescens 274 on a low-agar medium. J Bacteriol 177:987–91.

2. Partridge JD, Nhu NTQ, Dufour YS, Harshey RM. 2019. Escherichia coli Remodels the Chemotaxis Pathway for Swarming. mBio 10.

3. Dufour YS, Gillet S, Frankel NW, Weibel DB, Emonet T. 2016. Direct Correlation between Motile Behavior and Protein Abundance in Single Cells. PLoS Comput Biol 12:e1005041.

4. Stan Development Team. RStan: the R interface to Stan. 2019. http://mc-stan.org/

5. Bürkner, P. 2017. brms: An R Package for Bayesian Multilevel Models Using Stan. J. Stat. Soft. 10.18637/jss.v080.i01

6. Bürkner, P. 2018. Advanced Bayesian Multilevel Modeling with the R Package brms. The R Journal. 10:1, 395-4118.

7. R Core Team. 2018. R: A language and environment for statistical computing. R Foundation for Statistical Computing, Vienna, Austria. Available online at https://www.R-project.org/

7. Wickham, H. 2016. Package ‘ggplot2’: Elegant Graphics for Data Analysis. Springer-Verlag New York. doi:10.1093/bioinformatics/btr406

8. Kay, M. 2019. tidybayes: Tidy Data and Geoms for Bayesian Models. R Packag version 110. doi:10.5281/zenodo.1308151

